# Semi-continuous propagation of influenza A virus and its defective interfering particles: analyzing the dynamic competition to select candidates for antiviral therapy

**DOI:** 10.1101/2021.02.08.430251

**Authors:** Lars Pelz, Daniel Rüdiger, Tanya Dogra, Fadi G. Alnaji, Yvonne Genzel, Christopher B. Brooke, Sascha Y. Kupke, Udo Reichl

**Affiliations:** Max Planck Institute for Dynamics of Complex Technical Systems, Bioprocess Engineering, Magdeburg, Germany; University of Illinois at Urbana-Champaign, Department of Microbiology, Urbana, Illinois, USA; Otto-von-Guericke-University Magdeburg, Bioprocess Engineering, Magdeburg, Germany

**Keywords:** Influenza A virus, defective interfering particles, antiviral, next-generation sequencing, continuous virus production

## Abstract

Defective interfering particles (DIPs) of influenza A virus (IAV) are naturally occurring mutants that comprise an internal deletion in one of their eight viral RNA (vRNA) segments, rendering them propagation-incompetent. Upon co-infection with infectious standard virus (STV), DIPs interfere with STV replication through competitive inhibition. Thus, DIPs are proposed as potent antivirals for treatment of the influenza disease. To select corresponding candidates, we studied *de novo* generation of DIPs and propagation competition between different defective interfering (DI) vRNAs in a STV co-infection scenario in cell culture. A small-scale two-stage cultivation system that allows long-term semi-continuous propagation of IAV and its DIPs was used. Strong periodic oscillations in virus titers were observed due to the dynamic interaction of DIPs and STVs. Using next-generation sequencing, we detected a predominant formation and accumulation of DI vRNAs on the polymerase-encoding segments. Short DI vRNAs accumulated to higher fractions than longer ones, indicating a replication advantage. Yet, a sweet spot of fragment length was observed. Some DI vRNAs showed breaking points in a specific part of their bundling signal (belonging to the packaging signal), suggesting its dispensability for DI vRNA propagation. Over a total cultivation time of 21 days, several individual DI vRNAs accumulated to high fractions, while others decreased. Using reverse genetics for IAV, purely clonal DIPs derived from highly replicating DI vRNAs were generated. We confirm that these DIPs exhibit a superior *in vitro* interfering efficacy than DIPs derived from lowly accumulated DI vRNAs and suggest promising candidates for efficacious antiviral treatment.

**Importance:** Defective interfering particles (DIPs) emerge naturally during viral infection and typically show an internal deletion in the viral genome. Thus, DIPs are propagation-incompetent. Previous research suggests DIPs as potent antiviral compounds for many different virus families due to their ability to interfere with virus replication by competitive inhibition. For instance, the administration of influenza A virus (IAV) DIPs resulted in a rescue of mice from an otherwise lethal IAV dose. Moreover, no apparent toxic effects were observed when only DIPs were administered to mice and ferrets. IAV DIPs show antiviral activity against many different IAV strains, including pandemic and highly pathogenic avian strains, and even against non-homologous viruses, like SARS-CoV-2, by stimulation of innate immunity. Here, we used a cultivation/infection system, which exerted selection pressure toward accumulation of highly competitive IAV DIPs. These DIPs showed a superior interfering efficacy *in vitro*, and we suggest them for effective antiviral therapy.

## 1 Introduction

Yearly, on average 400,000 people globally die from an infection with seasonal influenza A virus (IAV) (1). Moreover, the potential emergence of pandemic strains is a major threat to public health (2). The most effective prevention of the influenza disease is vaccination with tri-or quadrivalent formulations, which provide protection against different influenza virus strains (3, 4). However, influenza vaccines have to be reformulated annually as a result of antigenic drifts (5). This is associated with a potential decrease in vaccine efficacy due to false predictions and a vaccine mismatch to circulating strains (6). Furthermore, antiviral drugs targeting the viral neuraminidase (oseltamivir, zanamivir) (7) or the viral endonuclease (baloxavir) (8) may also be used. Yet, circulating strains have already shown resistance against available antivirals (9-11). Therefore, the development of effective prophylactic and therapeutic treatment options is urgently needed.

One promising approach for antiviral therapy is the application of defective interfering particles (DIPs) (12-16). These naturally occurring viral mutants feature an internal deletion in one of their eight viral RNA (vRNA) segments, which renders them defective in virus replication. In addition, a new species of IAV DIPs that showed point mutations on segment (Seg) 7 vRNA was discovered recently (17). DIPs can only replicate in a co-infection with infectious standard virus (STV), which complements the respective defect in the replication of the DIPs. These viral mutants are believed to interfere by preferential and faster replication of the defective interfering (DI) vRNA in comparison to the full-length (FL) vRNA, thereby drawing away cellular and viral resources required for STV growth (18-20). Furthermore, interference was shown at the packaging step, as DI vRNAs can selectively outcompete FL vRNA packaging (21, 22). Notably, in mouse and ferret models, the administration of DIPs resulted in a pronounced antiviral effect against IAV infection (13, 14, 23-26). Furthermore, IAV DIPs also showed protection against heterologous interferon-sensitive respiratory viruses (27, 28), including SARS-CoV-2 (29), by the ability to stimulate innate immunity.

Recently, we established a two-stage bioreactor system for cell culture-based production of IAV (for vaccine manufacturing) (30), and of a prototypic, well-characterized DIP (“DI244” (23, 24, 27)) (31). Here, uninfected cells (first bioreactor) were continuously fed to a second bioreactor that contained virus-infected cells. However, in such a continuous culture, the co-infection of STVs and DIPs typically result in periodic oscillations of virus titers due to their dynamic interactions. Moreover, *de novo* generation and accumulation of numerous DI vRNAs was observed (30, 31).

In the present study, a simplified, semi-continuous setup was used to thoroughly investigate the generation and growth competition between DIPs during 21 days of IAV infection. Assuming that DIPs showing exceptional propagation also show high interfering efficacies, we anticipated identification of potent candidates for antiviral therapy. For detection and quantification of the different deletion junction on the IAV vRNA level, we used a recently published next-generation sequencing (NGS) framework (32). We observed a small subset of highly accumulated DI vRNAs after 21 days post infection (dpi), while other deletion junctions showed a pronounced decrease in their fractions in the same timeframe. To generate corresponding purely clonal DIPs harboring the promising candidate DI vRNAs, we used reverse genetics for IAV. Indeed, these DIPs displayed a superior *in vitro* interfering efficacy compared to DIPs derived from lowly replicating DI vRNAs indicating their potential for antiviral therapy.

## 2 Results

### 2.1 Semi-continuous production of IAV results in periodic oscillations of virus titers and strong accumulation of DIPs

In order to induce *de novo* generation and accumulation of DIPs, an IAV strain A/PR/8/34 of the subtype H1N1 (PR8, provided by Robert Koch Institute, Berlin, Germany) was propagated in a semi-continuous small-scale two-stage cultivation system (Fig. 1B). For infection, we used a seed virus that was depleted in DI vRNAs as shown by segment-specific reverse-transcription (RT-)PCR (Fig. 1A). Madin-Darby canine kidney (MDCK) cells growing in suspension culture (MDCK(sus)) were seeded into the cell seeding shake flask (CSS) and virus shake flasks (VS) at a viable cell concentration (VCC) of 0.6 × 10^6^ cells/mL and grown in batch mode to about 3.0 × 10^6^ cells/mL (−1.6 dpi) (Supplementary Fig. 1). Subsequently, for both shake flasks, a calculated volume of cell suspension was discarded and fresh medium was added at regular time intervals (both shake flasks not yet connected in series). This resulted in a residence time (RT) of 38.3 h for both vessels. Note that preliminary studies showed a steady state in the VCC for this RT (data not shown). Once the steady state was reached (Supplementary Fig. 1), cells in the VS were infected with PR8 at an MOI of 0.1. At 0.5 dpi, both vessels were connected in series and, from thereon, cells transferred semi-continuously from the CSS to the VS (V_2_). In addition, fresh medium was added to both shake flasks (V_1_ or V_3_) and virus harvest was taken (V_4_). The RT chosen was 38.3 h and 22.0 h for CSS and VS, respectively, as this previously resulted in pronounced titer fluctuations and strong accumulation of DIPs (31). Over the production time of 21 dpi, the steady state in the CSS was kept with an average VCC of 2.6 × 10^6^ cells/mL (SD of ±0.2 × 10^6^ cells/mL) (Supplementary Fig. 1).

**Figure 1:**
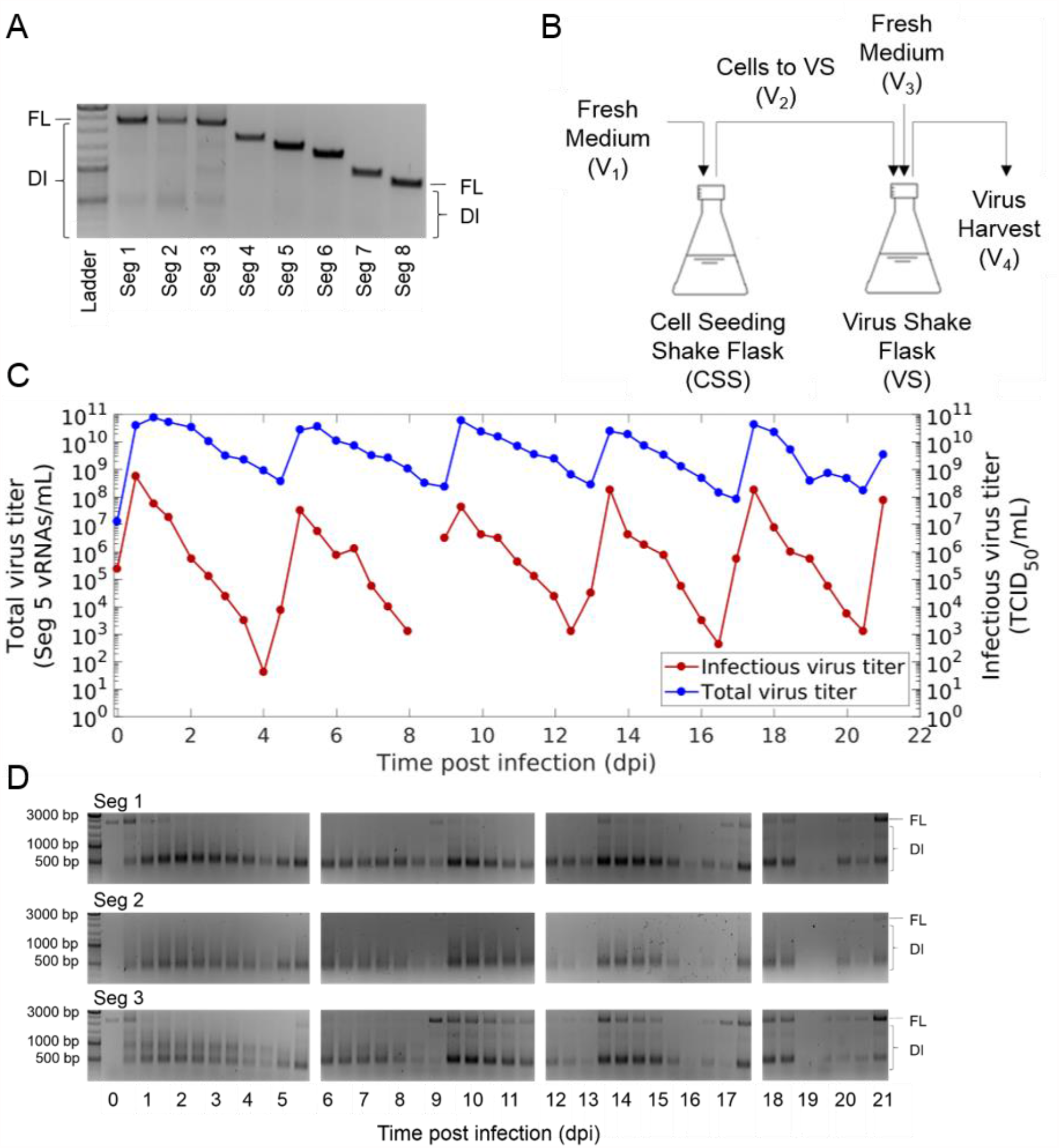
Semi-continuous propagation of influenza A virus and DIPs. (A) PR8 seed virus depleted in DI vRNAs was used for infection. Results of segment-specific RT-PCR for Seg 1 to 8 followed by agarose gel electrophoresis are shown. Signals corresponding to FL and DI vRNAs are indicated. Upper, middle and lower thick bands of the DNA ladder indicate 3000, 1000 and 500 bp, respectively. (B) Experimental setup of the small-scale two-stage cultivation system in shake flasks (scheme adapted from F. Tapia et al. (37)). MDCK(sus) cells were grown in the CSS and VS. After an initial batch and semi-continuous phase (CSS and VS not coupled), the cells in the VS were infected with the seed virus (A) at an MOI of 0.1. The semi-continuous production mode was initiated 0.5 dpi, where cells were transferred from the CSS into the VS (V_2_) at regular time intervals, while fresh medium was added (V_1_ or V_3_) and virus harvest was taken for monitoring (V_4_). (C) Periodic oscillations of total and infectious virus titers during the production. vRNA level of Seg 5 (indicating total virus particle concentration) was quantified by real-time RT-qPCR and infectious virus titer by TCID_50_ assay. D) Accumulation of DI vRNAs over the semi-continuous production time of 21 days. Results of the segment-specific RT-PCR are shown for Seg 1, 2, and 3. Signals corresponding to FL and DI vRNAs are indicated. Illustration includes results of one experiment.

Strong periodic oscillations in the infectious virus titers (quantified by TCID_50_ assay) and in the extracellular vRNA level of Seg 5 (quantified by real-time RT-qPCR) were observed in the VS (Fig. 1C). The extracellular vRNA level of Seg 5 was taken as a measure of the total virus concentration. DI vRNAs are mostly located on polymerase-encoding segments (20, 33-36), so the occurrence of DIPs should not affect the detection of Seg 5 vRNA. Shortly after infection at 0.5 dpi, a maximum infectious virus titer of 5.6 × 10^8^ TCID_50_/mL was reached. Here, high concentrations of STV (complying with a high MOI) increased the chance for co-infections with DIPs. Thus, a strong DIP propagation likely occurred early in cultivation, impeding STV propagation. Therefore, infectious virus titers decreased from 0.5 dpi onwards. Eventually, the declining infectious virus titers led to fewer co-infections. Thus, DIP replication decreased, and the total virus particle concentration dropped as well. Additionally, DIPs were out-diluted because of the semi-continuous feeding strategy. Then, at a low infectious virus concentration (complying with a low MOI condition, ∼4.0 dpi), the chance of DIP co-infections was supposedly significantly reduced. In these conditions, STVs could accumulate again as indicated by increasing virus titers toward 5 dpi. In the following, further periodic oscillations in virus titers occurred based on the DIP/STV interaction described above.

The dynamics in virus titers were well in agreement with results of the segment-specific RT-PCR (indicating FL and DI vRNAs) (Fig. 1D). A rapid accumulation of DI vRNAs occurred already at 0.5 dpi. Furthermore, the FL vRNA signal gradually dropped between 1 dpi and 2.5 dpi, suggesting the preferential production of DI vRNAs. Subsequently, DI vRNA replication decreased and DI vRNAs were washed out, as indicated by weaker band intensities of DI vRNAs (e.g. at 8.5 dpi). Next, in agreement with the increase of infectious viral titers (STVs), FL vRNA bands were visible again (e.g. at 9 dpi). Moreover, agarose gels indicated the presence of DI vRNA bands at the end of cultivation that may have been already present in the seed virus, suggesting that some DI vRNAs were preserved. In addition, weak DI vRNA bands as well as undefined, blurred bands emerged during the course of IAV replication, suggesting the formation and accumulation of *de novo* generated DI vRNAs.

In summary, the semi-continuous production of IAV using a seed virus depleted in DI vRNAs led to the accumulation of DIPs. Thus, strong periodic oscillations in the total concentration of virions and infectious virus titers were observed due to the dynamic interaction of STVs and DIPs. Moreover, in the course of production, DIPs were exposed to high and low MOI conditions that have likely resulted in alternating selection pressures, suitable for potential selection toward accumulation of highly interfering DIPs.

### 2.2 Next-generation sequencing results indicate predominant *de novo* formation and accumulation of deletion junctions on polymerase-encoding segments

Segment-specific RT-PCR does not enable the detection and quantification of individual deletion junctions. Therefore, to study the diversity of DI vRNAs generated during semi-continuous IAV propagation, samples were subjected to Illumina-based NGS and processed by a bioinformatics pipeline (32). Doing so, sequences of vRNAs from the produced progeny virions were obtained. Reads comprising a deletion junction (DI vRNA reads) do not align to the corresponding reference genome. These NGS reads were processed by the ViReMa algorithm to identify the position of individual deletion junctions (38).

The highest variation (i.e., number of different deletion junction) was found on the polymerase-encoding segments 1–3, which encode for the polymerase basic protein 2 (PB2) and 1 (PB1), and polymerase acidic protein (PA), respectively (Fig. 2A). Fig. 2B shows the fraction of all deletion junctions located on a genome segment over time. Here, polymerase-encoding segments showed the highest fraction. In contrast, deletion junctions of non-polymerase-encoding segments showed a significantly lower fraction, which increased slightly toward the end of cultivation but always remained below 2%. As non-polymerase segment deletion junctions occurred only in negligible numbers, they were not considered any further in subsequent analyses.

**Figure 2:**
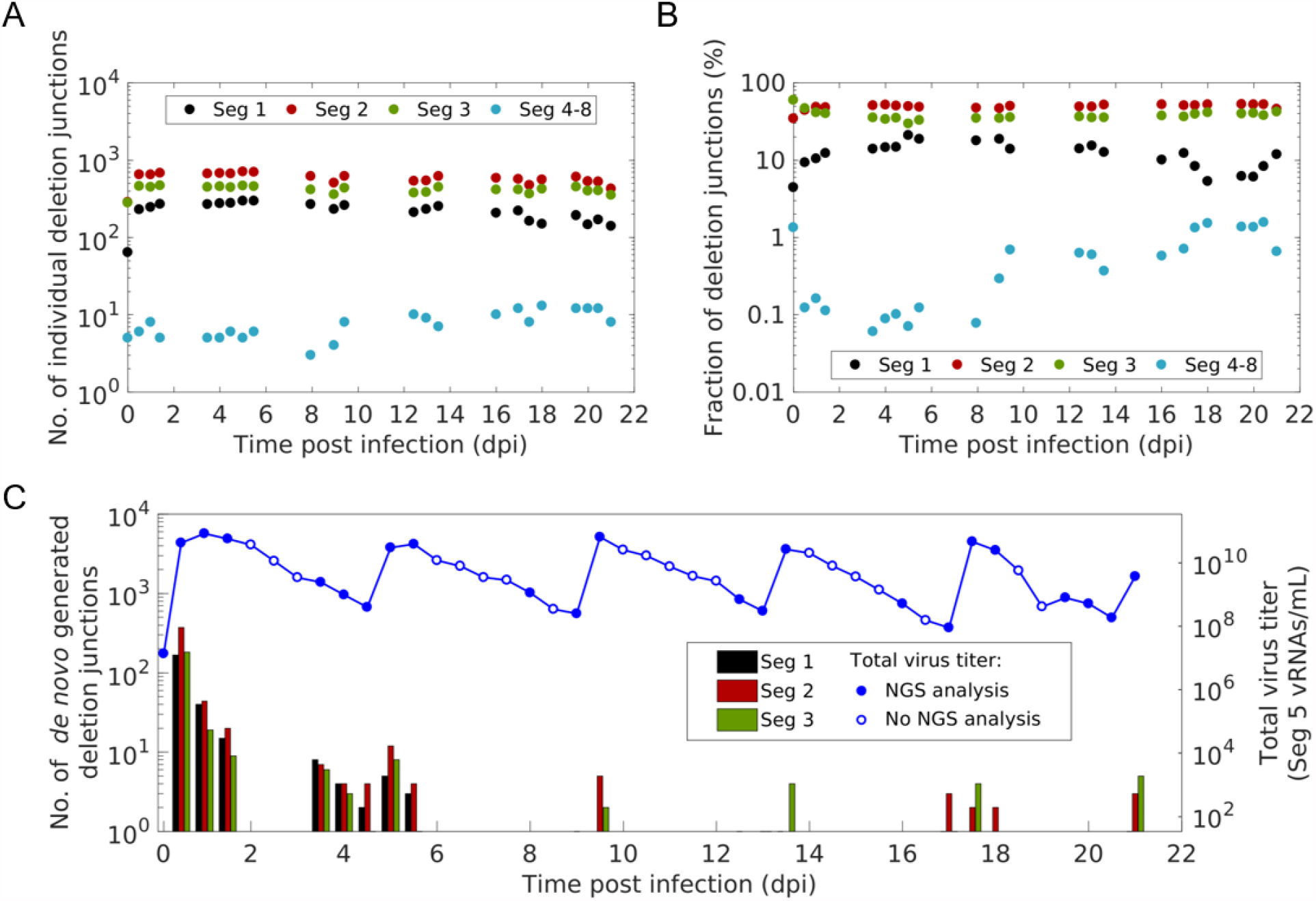
Diversity, distribution and *de novo* generation of deletion junctions during semi-continuous propagation of IAV. Deletion junctions were identified by Illumina-based NGS and subsequent analysis via the ViReMa algorithm (32). (A) Number of different deletion junctions located on the respective genome segment(s). (B) Fraction of all deletion junctions located on the respective genome segment(s). This fraction describes the ratio of the total number of detected deletion junctions for one segment to the total number of deletion junctions on all genome segments. (C) *De novo* formation of deletion junctions. vRNA level of Seg 5 (indicating total virus particle concentration, as shown in Fig. 1C) was quantified by real-time RT-qPCR. Samples not analyzed by NGS are indicated by blank dots. Illustration includes results of one experiment.

Next, we investigated the *de novo* formation of DI vRNAs over the course of the cultivation. Fig. 2C shows, at specific time intervals, the number of *de novo* generated deletion junctions. *De novo* formation occurred mainly on the polymerase-encoding segments. Interestingly, most *de novo* formations occurred within the first 1.5 dpi. In addition, a considerable number of *de novo* DI vRNAs was detected between 3.5–5.5 dpi. However, *de novo* formation was significantly lower at later time points. Moreover, an increase in the number of new deletions was highly correlated with an increase in the total virus particle concentration (indicated by the vRNA level of Seg 5) (Fig. 2C). This is consistent with a fast STV replication, and thus, likely with a higher occurrence of the *de novo* formation of DI vRNAs due to the error-prone nature in the replication of the IAV RNA-dependent RNA polymerase.

In sum, our results show that DI vRNAs are predominantly *de novo* formed and accumulated on the polymerase-encoding segments during semi-continuous IAV infection.

### 2.3 Short DI vRNAs tend to accumulate to higher fractions than longer ones; yet, intermediate length sweet spots were observed as well

It was reported that DI vRNAs accumulation surpasses that of FL vRNAs due to their shorter length resulting in a supposedly faster replication (12, 20). Therefore, we speculated that shorter DI vRNAs may also accumulate to higher abundances than longer DI vRNAs. Fig. 3 shows the fraction of all individual deletion junctions and their corresponding DI vRNA length. Indeed, a bias toward accumulation of shorter DI vRNAs was observed with short DI vRNAs showing overall higher fractions than longer ones during semi-continuous IAV production (Fig. 3). However, highest fractions were not found for the shortest DI vRNAs. It rather appeared that highest fractions were found at a length sweet spot. To visualize this sweet spot, we fitted a normal distribution function to the DI vRNA length (Supplementary Fig. 2), and plotted the resulting mean as a dashed vertical line (Fig. 3). Over the whole cultivation, the mean DI vRNA length ranged between 366– 414 nt, 425–534 nt, and 434–557 nt for Seg 1, 2, and 3, respectively.

**Figure 3:**
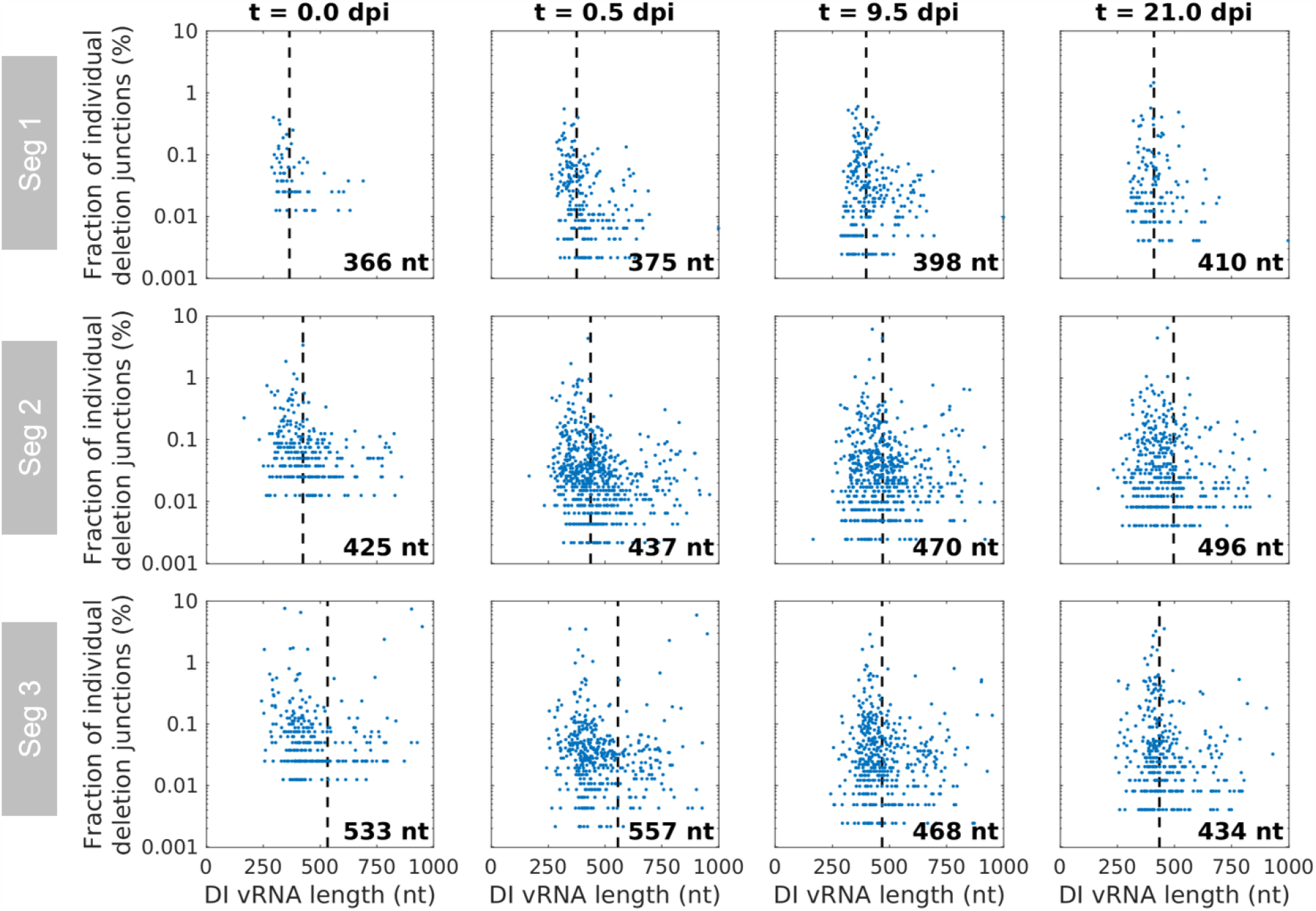
Dependency of the length of DI vRNAs on their accumulation during semi-continuous propagation of IAV. Deletion junctions were identified by Illumina-based NGS and subsequently analyzed via the ViReMa algorithm (32). Fractions of individual deletion junctions were calculated based on the ratio of the number of NGS reads of one individual deletion junction to the number of NGS reads of all deletion junctions located on all eight segments. Means of DI vRNA length (calculated by fitting a normal distribution function) are indicated by dashed vertical lines and corresponding lengths are shown. Representative time points are illustrated. Illustration includes results of one experiment.

Moreover, a few larger DI vRNAs (comprising a sequence length of up to 1000 nt) accumulated to high fractions, suggesting that the sequence and the position of the deletion junction may be another factor to consider for replication of a DIP (Fig. 3). Note that Fig. 3 only shows DI vRNAs up to 1000 nt in length, although we also detected very long DI vRNAs (>2000 nt) (Supplementary Fig. 3). These DI vRNAs with very short deletions may either not result in a defective vRNA, comprise two deletions or represent technical artifacts. Due to their unknown origin and function and a lack of description in the literature, defective vRNAs larger than 85 % of its respective FL length were excluded from analysis in this work (shown in Supplementary Fig. 3).

Taken together, shorter DI vRNAs showed an overall stronger accumulation compared to longer DI vRNAs. However, highest fractions were found at a sweet spot, indicative for an optimal length for efficient DI vRNA replication and spreading.

### 2.4 The incorporation signal but not the entire bundling signal appears to be required for propagation of DIPs

We next examined the position of the breaking points of DI vRNAs. Fig. 4 illustrates the position of individual deletion junctions, as indicated by the number of retained nucleotides prior (DI vRNA 3’ length) and after (DI vRNA 5’ length) the deletion junction site. In the course of the semi-continuous cultivation, breaking points were mostly located in proximity to both ends of vRNA (Fig. 4). This finding is in line with our observation of the predominant accumulation of short DI vRNAs (Fig. 3.). We also observed highly abundant medium-sized DI vRNAs on Seg 3 in the seed virus (0.0 dpi). Yet, the fraction of DI vRNAs carrying these deletions decreased, or even disappeared toward the end of cultivation (Fig. 3). Again, this indicates that shorter DI vRNAs replicate faster and may outcompete longer ones. Additionally, the 3’ length of the DI vRNA did largely not correlate with the 5’ length, suggesting that deletion junctions are not preferably symmetrical.

**Figure 4:**
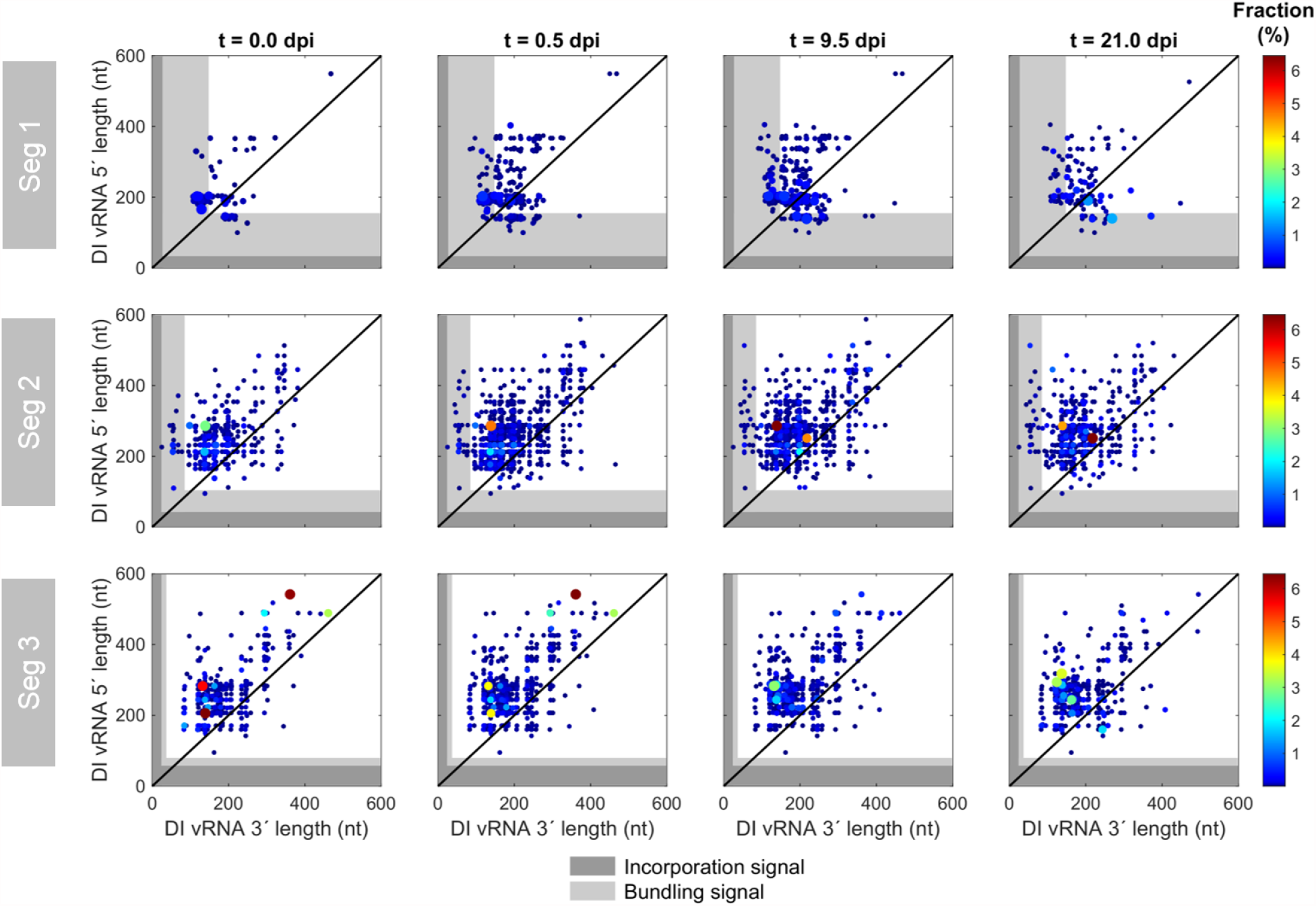
Deletion junction sites of DI vRNAs during semi-continuous propagation of IAV. Deletion junctions were identified by Illumina-based NGS and subsequently analyzed via the ViReMa algorithm (32). DI vRNA 3’ and 5’ length indicate the number of retained nucleotides prior and after the deletion junction, respectively, at the corresponding vRNA ends. The packaging signal is indicated as grey areas, which is divided into the incorporation signal (dark grey area) and bundling signal (light grey area). Representative time points are illustrated. The color code from red to blue shown on the right denotes the fraction of individual deletion junction, which was calculated based on the ratio of the number of NGS reads of one individual deletion junction to the number of NGS reads of all deletion junctions located on all eight segments. Additionally, the circle radii increase with higher fractions. The diagonal black line indicates an equal DI vRNA 3’ and 5’length. Illustration includes results of one experiment.

While the lengths of the 3’ and 5’ ends ranged from below 100 nt to over 500 nt, specific minimum lengths were retained in the DI vRNAs (Fig. 4). We then asked whether the complete packaging signal (situated at the terminal ends of vRNA), which is important for organized packaging into progeny virions (39), was unaffected by deletions. A small percentage of breaking points was located in the packaging signal (on Seg 1 and 2); yet, the majority of the deletion junction sites were located outside of it, which is in line with the observation of a sweet spot in DI vRNA length (Fig. 3). For a more thorough investigation of deletion junctions in the packaging signal, we highlighted the positions of the incorporation signal (non-coding region (NCR), including the promoter region) and the bundling signal (terminal ends of coding region) (40). The incorporation signal was reported to lead the packaging of the vRNA in which the signal is found. The second part of the packaging signal is the bundling signal, which confers the selective packaging of all the eight different segments together (40). We checked which part of the sequence at both ends were retained to infer a minimum sequence length for functional replication and packaging of the truncated vRNAs, assuming that only propagation-competent DI vRNAs can be detected. No deletion junctions in the incorporation signal for the polymerase-encoding segments as well as for Seg 4–8 were identified (Fig. 4, Supplementary Fig. 4, respectively). Therefore, we suggest that the preservation of the entire incorporation signal is crucial for the propagation of DIPs

Interestingly, deletion junctions in the bundling signal (on Seg 1 and 2) could be detected, indicating that the entire bundling signal of these segments is most likely not required for propagation of DIPs. In particular, clusters of DI vRNA breaking points in the bundling signal were stable and present over the complete course of the semi-continuous cultivation. In contrast, Seg 3 did not show any breaking points in both signals. We found a minimum sequence length of 84 nt (3’ end) and 100 nt (5’ end), 25 nt and 95 nt, and 82 nt and 95 nt for Seg 1, 2, and 3, respectively. Supplementary Fig. 4 shows the position of deletion junction sites in Seg 4–8. Notably, although only very few individual deletion junctions were detected, breaking points were found in the bundling signal on Seg 6 (3’ end), Seg 7 (both ends), and Seg 8 (5’ end) as well.

In summary, our results indicate that the complete incorporation signal is crucial for propagation of DIPs. Yet, only a part of the bundling signal in Seg 1 and 2 seem to be required for DIP spreading.

### 2.5 Dynamic competition in propagation between DI vRNAs leads to selection toward accumulation of highly interfering DIPs

In order to elucidate whether the various DI vRNAs show differences in their propagation, we next studied the composition of deletion junctions over cultivation time. More specifically, we determined the fraction of each individual deletion junction over time. Fig. 5A shows these fractions and highlights the top five deletion junctions that showed the highest gain or largest loss in their fraction from the seed virus (0 dpi) to the end of cultivation (21 dpi). Likewise, the top five gains of *de novo* formed DI vRNAs are indicated. Interestingly, differences between gains and losses were very pronounced, with a decreasing fraction of the top five losses, while the top five gains (including *de novo*) showed a strong accumulation. These trends were most prominent for Seg 3. Of note is also one deletion junction on Seg 2 that was present at a very high fraction in the seed virus and throughout the whole cultivation. Furthermore, pronounced shifts in the composition of deletion junctions were found for 9–9.5 and 17–17.5 dpi, at best visible for Seg 3. The occurrence of DI vRNAs that accumulate faster and achieve higher fractions than other DI vRNAs suggests that there was a dynamic competition in the propagation between individual DI vRNAs.

**Figure 5:**
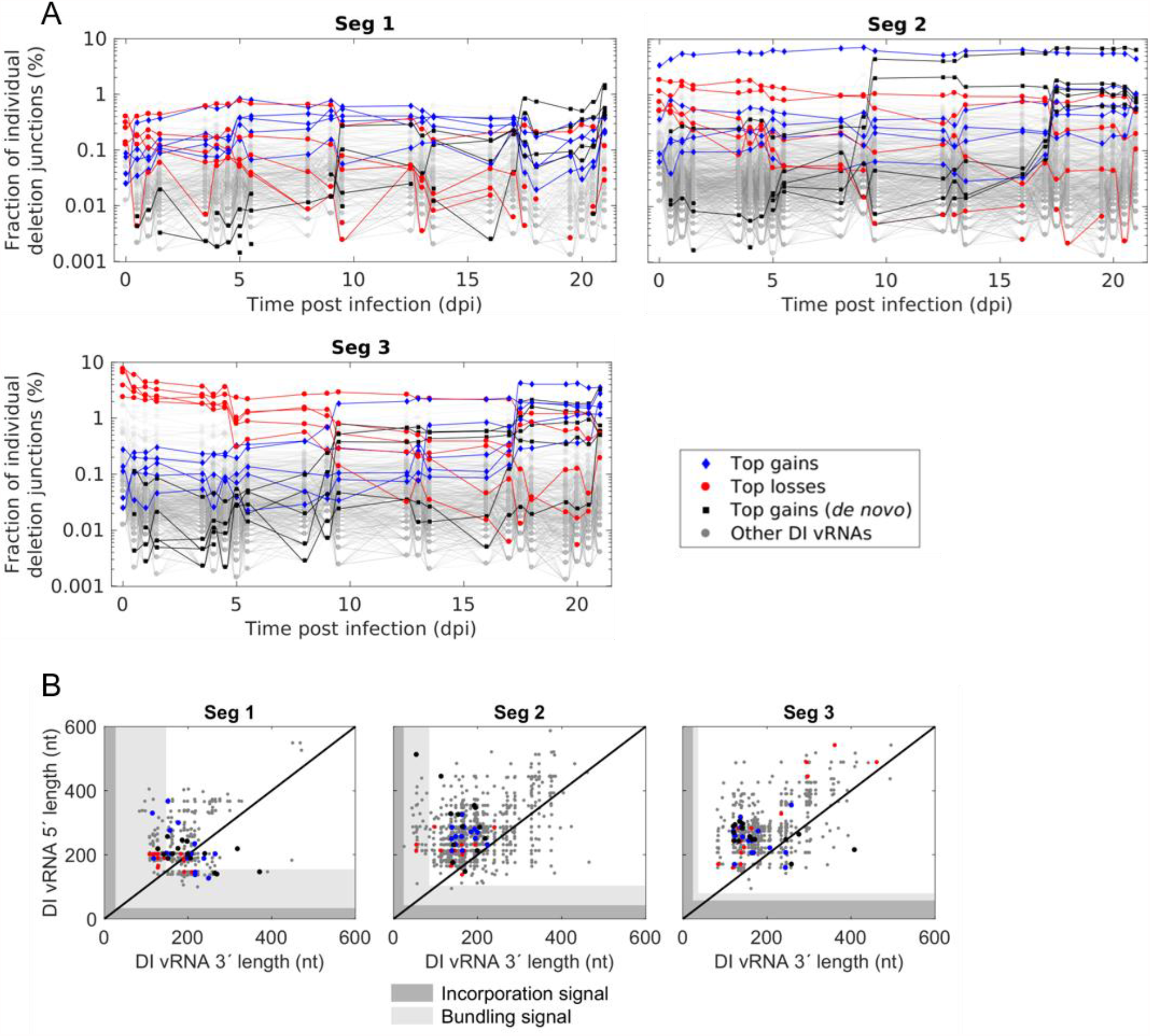
Propagation of DI vRNAs showing highest gains or losses in their fractions during semi-continuous propagation of IAV. Deletion junctions were identified by Illumina-based NGS and subsequently analyzed via the ViReMa algorithm (32). Fractions of individual deletion junctions were calculated based on the ratio of the number of NGS reads of one individual deletion junction to the number of NGS reads of all deletion junctions located on all eight segments. (A) Fraction of individual DI vRNAs belonging to the group of top five gains, losses, and gains (*de novo*) of the fraction over cultivation time. Top gains (*de novo*) indicate newly formed deletion junctions with the highest fraction at the end of cultivation. (B) Deletion junction position of the top 15 gains, losses, and gains (*de novo*). DI vRNA 3’ and 5’ length indicate the number of retained nucleotides prior and after the deletion junction, respectively, at the corresponding vRNA ends. The packaging signal is indicated as grey areas, which is divided into the incorporation signal (dark grey area) and bundling signal (light grey area). Illustration includes results of one experiment.

Moreover, we examined whether top gains (including *de novo*) and losses show differences in the deletion junction position (Fig. 5B). To obtain a better overview, we expanded the number of the top candidates in each category to 15. However, it appeared that no clear differences between the groups were present. For both gains and losses, few deletion junction sites were located in the bundling signal for Seg 1 and 2 (although most were found outside (Fig. 5B)), but none for Seg 3. Therefore, even for competitive DIPs (which require an efficient packaging process), we found a shorter packaging signal compared to the FL vRNA on Seg 1 (both ends) and on Seg 2 (3’ end). Please also note a few DI vRNAs (belonging to the top 15 losses) on Seg 3 showing a medium-sized DI vRNA length (∼900 nt) (Fig. 5B, upper right corner), which is in line with our observation that long DI vRNAs accumulate to low fractions (Fig. 3). Yet, we also found two top 15 gains (*de novo*) on Seg 2 with a very long DI vRNA (1905 nt and 1628 nt) (Supplementary Fig. 5). This finding might suggest that not only the sequence length but also the breaking point position and probably further unknown regulatory effects are crucial for the efficient propagation of DI vRNAs.

In order to test the hypothesis, that fast-propagating DI vRNAs show a higher interfering efficacy than slow-propagating ones, we reconstituted the corresponding DIPs and tested them in an *in vitro* interference assay. More specifically, we rescued purely clonal DIPs (in the absence of STV) harboring either the top gain, top loss, or top gain (*de novo*) DI vRNA of Seg 1 (Supplementary Table 5) using a modified reverse genetics system for IAV DIPs (25, 41). Next, we propagated these selected DIPs in genetically engineered MDCK-PB2(sus) cells, expressing PB2 (Supplementary Fig. 6), to allow multiplication of these DIPs (harboring a deletion in Seg 1) without STV through complementation. Almost complete absence of contamination with other DI vRNAs was confirmed by results of segment-specific RT-PCR (Supplementary Fig. 7).

In the *in vitro* interference assay (Fig. 6), adherent MDCK cells (MDCK(adh) cells were infected with STV at an MOI of 0.01 and co-infected with the different DIPs to evaluate the inhibition of STV replication compared to infection with STV alone (NC). Here, the DIP input for the interference assay was normalized through dilution based on the concentration of DIPs to ensure a direct comparison between the DIPs (Supplementary Table 7). In addition, we compared the interfering efficacy to a prototypic, well-characterized DIP named DI244 (23-25, 27). Indeed, the DIPs derived from the top gains (including *de novo*) showed the highest interfering efficacy. The top gain (*de novo*) DIP reduced the infectious virus release by more than five orders of magnitudes, the top gain by five logs, while top loss and DI244 showed a reduction of only four orders of magnitudes. Reduction of the total virus particle release, indicated by the HA titer, showed a similar trend (Fig. 6B).

**Figure 6:**
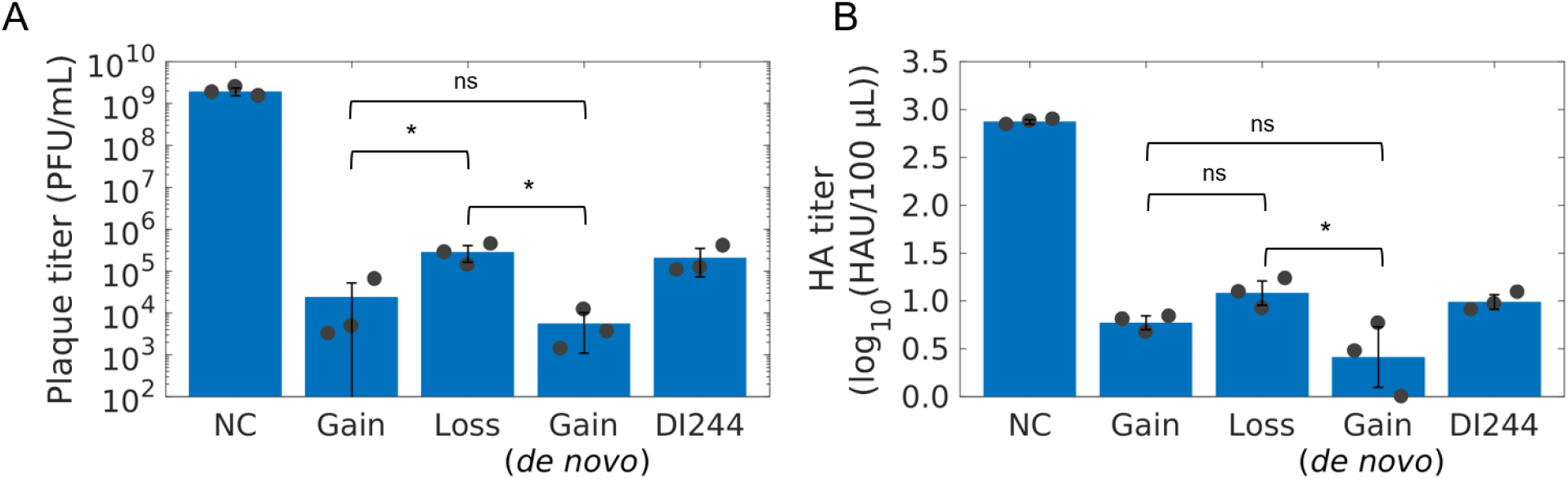
Interfering efficacy of DIPs derived from DI vRNAs showing highest gain, loss or gain (*de novo*) in their fraction during semi-continuous propagation of IAV. Purely clonal DIPs containing a deletion in Seg 1 (derived from DI vRNAs showing top gains and top loss) were generated using a modified reverse genetics methodology for reconstitution of purely clonal IAV DIPs (41). Next, DIPs were multiplied in genetically engineered MDCK-PB2(sus) cells (expressing PB2) in shake flask at a multiplicity of DIP (MODIP) E-2 (Supplementary Fig. 6). (A and B) Interference assay. MDCK(adh) cells were infected with STV only at an MOI of 0.01 (NC) or co-infected with the corresponding DIP resulting in a DIP/STV ratio of 3224 (number of DIPs: 4.25 × 10^8^ virions, number of STVs: 1.32 × 10^5^ virions, derived from the HA titer, Supplementary Table 7). For comparison, DI244, a prototypic, well-characterized DIP (23-25, 27), was used. (A) Infectious virus release, quantified by plaque assay. (B) Total virus particle release, measured by HA assay. Illustration includes results of three independent experiments (n = 3). Error bars indicates standard deviation. One-way ANOVA followed by Tukey’s multiple comparison test (* p < 0.05; ns p > 0.05, not significant) was used to determine significance.

In summary, our results indicate that the semi-continuous propagation of IAV led to a dynamic competition in propagation between different DI vRNAs. We demonstrate that DI vRNAs showing the highest increase in the fraction over the cultivation period result in the formation of DIPs that show a superior interfering efficacy compared to DIPs containing slowly propagating DI vRNAs. These DIPs are, thus, promising candidates for antiviral therapy.

## 3 Discussion

IAV DIPs have been proposed as an effective antiviral agent for the influenza disease. In this study, we investigated the *de novo* generation and the competition in the growth between a diversity of DIPs during long-term semi-continuous IAV infection in order to identify strong candidates for antiviral therapy. In general, DIPs and STVs are in a competition for cellular and viral resources in a co-infection scenario (19, 42). Due to replication advantage of DIPs, suppression of and interference with STV replication occurs (18-20). Moreover, it was shown that DIPs interfere with STV propagation at the packaging step, as preferential incorporation of DI vRNAs over FL vRNAs was observed (21, 22). We, thus, hypothesized that DI vRNAs showing the strongest accumulation during long-term co-infection possess the highest interference efficacy with STV replication. In our experiments, a small subset of individual DI vRNAs was observed that showed a pronounced accumulation while the fractions of some other DI vRNAs strongly decreased (Fig. 5A). Next, DIPs harboring the most competitive DI vRNAs on Seg 1 were generated and we show that these DIPs exhibit a higher interfering efficacy than slowly propagating ones (Fig. 6). Strikingly, the interfering efficacy was also higher in comparison to DI244, a prototypic and well-characterized DIP (23-25), suggesting a huge potential of these candidates for antiviral treatment.

Our data show that the most competitive DI vRNAs are derived from the polymerase-encoding segments. Further, we found the highest variation, accumulation and *de novo* formation of DI vRNAs on these segments (Fig. 2A, Fig. 2B, Fig. 2C, respectively). This confirms previous studies, which showed that DI vRNAs are predominantly found on Seg 1, 2, and 3 (20, 33-36). In agreement with this, a bias toward the emergence of DI vRNAs on the polymerase-encoding segments was observed during production of IAV over 17 d in a fully-continuous two-stage bioreactor system (30). Mathematical modeling of the intracellular replication during STV and DIP co-infection also suggested that DI vRNAs located on the polymerase-encoding segments are more competitive than DI vRNAs on other segments (18). In particular, they even yielded high progeny numbers in less advantageous infections scenarios, i.e., when STV co-infection was delayed by several hours. Accordingly, in *in vivo* studies, DI244 containing a deletion on Seg 1, or DI vRNAs carrying deletion in Seg 1, 2 or 3 showed a pronounced antiviral effect upon administration against IAV replication in mice and in ferrets (14, 23-25)

Our results show that short DI vRNAs tended to accumulate to higher fractions than longer DI vRNAs (Fig. 3). In general, it is believed that the shorter length of DI vRNAs (in comparison to FL vRNAs) leads to a replication advantage (18-20), which supports our findings. However, our observation of a length sweet spot also indicates an optimal length for DI vRNA replication and spreading (Fig. 3). In agreement with this, the highly potent DI244 (395 nt) shows a similar DI vRNA length (Fig. 3) (23-25, 27). Other studies reported a similar mean DI vRNA length of 400– 500 nt for Seg 1, 2, and 3 (12, 34). For DIPs originating from clinical isolates, a similar mean DI vRNA length of 377 nt (Seg 1), 390 nt (Seg 2), 490 nt (Seg 3) was found (36). Another investigation confirmed the finding of a replication advantage toward shorter vRNAs, but additionally suggested that the sequence (UTRs and coding region) may also have an influence on vRNA competition (42). This is consistent with a previous work proposing that not only the length, but also the sequence (or the deletion junction position) may drive the replication advantage of DI vRNAs (20). This may explain our observation of few larger DI vRNAs up to 1000 nt, which accumulated to high fractions (Fig. 3). In addition, two very long DI vRNAs (1905 nt and 1628 nt) were included in the top 15 gains (*de novo*) DI vRNAs (Supplementary Fig. 5). In further agreement, the *in vivo* interfering efficacy of three clonal DIPs containing DI vRNAs with a similar length (but diverse deletion junctions), differed significantly from each other (24). We found no clear patterns between the deletion junction positions of top gains (including *de novo*) and losses (Fig. 5B). These results may support the hypothesis that not only the DI vRNA length, but also the deletion junction site and further unknown regulatory effects are decisive factors for a competitive DI vRNAs.

Packaging of progeny virions of IAV is a complex process. The leading packaging model postulates a selective packaging of the eight different vRNA genome segments (43, 44). Decisive for correct and efficient packaging is a special vRNA sequence (packaging signal), which was discovered using reverse genetics approaches (39). As DIPs comprise a truncated segment, this packaging signal might equally be affected. However, it was suggested that this packaging signal in the shortened segment typically remains intact (12, 43). The packaging signal is divided into two parts, the incorporation signal (NCR, including promoter region) and the bundling signal (located at the terminal ends of the coding region). Our results show that the incorporation signal is crucial for DIP propagation as it was unaffected by deletions (Fig. 4). However, several deletion junction sites were located in the bundling signal (Fig. 4, Fig. 5B, Supplementary Fig. 4). Therefore, we suggest that DIPs require only a part of the bundling signal for efficient replication and spreading. This finding does not agree with a previous study, which implied that the entire packaging signal of is crucial for DI vRNA stability (45) and for high interference (46).

Previous works showed that DI vRNAs interfere with FL vRNAs at the packaging process, by selectively suppressing the packaging either of the parent segment (21, 22) or the FL vRNA of another genome segment (47). However, one recent study showed the opposite, in which DI vRNA were packaged less frequently than FL vRNA (previously posted on a preprint server (48)). Furthermore, differences in the packaging rates were found between individual DI vRNAs (21, 46). Thus, the highly abundant DI vRNAs found in the present study may have an advantage in the entire propagation process over others, including both replication and packaging. Yet, further in-depth studies are required to better characterize the interference of DI vRNAs at the virus assembly step.

Taken together, we show that DIPs containing DI vRNAs with a superior propagation rate also show a superior capacity to interfere with STV replication. These DIPs are very interesting candidates for antiviral treatment. The highly competitive DI vRNAs are predominantly located on the polymerase-encoding segments, display an optimal DI vRNA length, and conserve the incorporation signal, but do not require the entire bundling signal. In addition, yet unidentified sequence motifs certainly also play an additional role during DI vRNA propagation. Due to the complex features of highly competitive DIPs, the best candidates for antiviral therapy are probably challenging to design *in silico*. Thus, evolution studies are a more convenient screening tool as shown for DIPs of other virus families (49-51).

## 4 Materials and Methods

### 4.1 Cells and viruses

MDCK(adh) cells (ECACC, No. 84121903) were adapted in previous works to grow in suspension culture (52), and then in chemically defined Xeno^™^ medium (53), in this work referred to as MDCK(sus) cells. Further, this cell line was engineered to stably express the PB2 for the production of purely clonal DIPs harboring a DI vRNA in Seg 1 (25, 41) and is denoted as MDCK-PB2(sus). Cultivation of both cell lines was conducted in shake flasks at a working volume of 50 mL (125 mL baffled Erlenmeyer Flask, Thermo Fisher Scientific, 4116-0125) using an orbital shaker (Multitron Pro, Infors HT; 50 mm shaking orbit) at 185 rpm and 37°C in a 5% CO_2_ environment. The medium was supplemented with 8 mM glutamine. For MDCK-PB2(sus) cells, puromycin (Thermo Fisher Scientific, #A1113803) was added at a final concentration of 0.5 μg/mL. Quantification of VCC, viability and diameter were performed using a cell counter (Vi-Cell™ XR, Beckman coulter, #731050). MDCK(adh) cells were maintained in Glasgow minimum essential medium (GMEM, Thermo Fisher Scientific, #221000093) supplemented with 10% fetal bovine serum and 1% peptone at 37°C and 5% CO_2_. The corresponding adherent MDCK cell line that stably express PB2 (MDCK-PB2(adh)) (41) was maintained in the presence of 1.5 μg/mL puromycin. Adherent PB2-expressing HEK-293T (HEK-293T-PB2) cells (41) were maintained in Dulbecco’s Modified Eagle Medium (DMEM, Gibco, #41966029) supplemented with 10% fetal bovine serum, 1% penicillin streptomycin (10,000 units/mL penicillin and 10,000 µg/mL streptomycin, Gibco, #15140122) and 1 µg/mL puromycin at 37°C and 5% CO2.

For virus infection during semi-continuous cultivation, PR8 (provided by Robert Koch Institute, Berlin, Germany) was used (53). The strain was adapted to MDCK(sus) cells and depletion of DI vRNAs was carried out over five passages at a very low MOI of 10^−5^. For the interference assay in MDCK(adh) cells, the same PR8 strain, but adapted to adherent MDCK cells was used. In addition, we generated candidate DIPs containing a deletion in Seg 1 using reverse genetics as described in chapter 4.6.

### 4.2 Small-scale two-stage cultivation system for semi-continuous STV/DIP propagation

For the semi-continuous propagation of PR8, a two-stage cultivation system was used, which consisted of two baffled shake flasks (250 mL baffled Erlenmeyer Flask, Thermo Fisher Scientific, 4116-0250) connected in series (Fig. 1B). The CSS and the VS were operated at a working volume of 90.00 mL and 77.52 mL, respectively. MDCK(sus) cells in exponential growth phase were seeded at a VCC of 0.6 × 10^6^ cells/mL and were cultivated in batch mode at 185 rpm, and 37°C in a 5% CO_2_ environment for 2 days. When VCC reached approximately 3.0 × 10^6^ cells/mL, at -1.6 dpi, a calculated volume of cell suspension was harvested every 12 h, while pre-warmed fresh medium was added manually to obtain a RT (inverse of the dilution rate) of 38.3 h (both, CSS and VS). Please note that both shake flasks were not yet connected in series. After steady state was achieved, the cells in the VS were infected with PR8 at an MOI of 0.1 and trypsin (Thermo Fisher Scientific, #27250-018) was added (final activity of 20 U/mL). At 12 hpi, semi-continuous production was started by transferring cells from the CSS to the VS (V_2_). Furthermore, virus was harvested (V_4_) and both shake flasks were filled up with pre-warmed fresh medium (V_1_ or V_3_) to obtain a RT of 38.3 h and 22.0 h for CSS and VS, respectively. Of importance is that the fresh medium, which was added to the VS, contained 60 U/mL trypsin to reach 20 U/mL in the VS. The respective transferred volumes are indicated (Equation 1–4). The RT of 22.0 h for the VS was chosen as previously published data showed a pronounced DIP/STV replication dynamic (31). In addition, samples were taken from the virus harvest at every volume transfer for analysis. Cell-free supernatants (300×g, 4°C, 5 min) were stored at -80°C for further analysis.

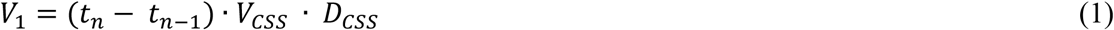

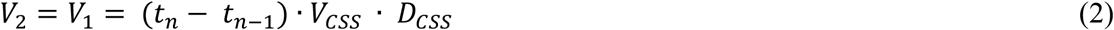

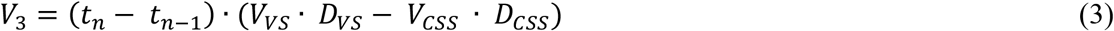

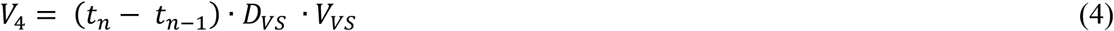

Where t_n_ denotes the sample time point, and t_n-1_ the previous sample time point. V_CSS_ is the volume of the CSS, V_VS_ is the volume of VS, D_CSS_ is the dilution rate of CSS, D_VS_ is the dilution rate of VS. V_CSS_, D_CSS_, D_VS_ were predefined as mentioned above. V_3_ was set as 0.5 × V_2_ to ensure a sufficient volume of fresh medium in the VS. This assumption was applied to calculate the volume of V_Vs_.

### 4.3 Virus quantification

Quantification of the infectious virus titer was performed by TCID_50_ assay as described previously (54) with a measurement error of ± 0.3 log_10_ (55). The active DIP titer (required for calculation of an MODIP for production of candidate DIPs in shake flasks, chapter 4.6.2) was quantified by plaque assay using MDCK-PB2(adh) cells (measurement error of ≤ 43.8%) (25). To determine the infectious virus titers in the interference assay (chapter 4.7), MDCK(adh) cells were deployed in the same plaque assay (25). In addition, an HA assay was used to quantify the total number of virions in the supernatant with a measurement error of ±0.15 log_10_(HAU/100 µL) (56). Concentration of DIPs (c_DIP_) or concentration of STVs (c_STV_) were derived from the HA titer and determined according to Equation 5, where c_RBC_ denotes the concentration of red blood cells (2.0 × 10^7^ cells/mL).

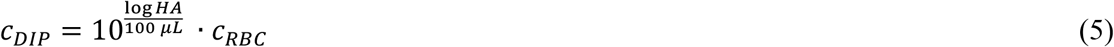

### 4.4 PCR measurements

Genomic vRNA in progeny virions were examined using PCR. In brief, isolation of vRNA from 150 µL cell-free supernatants was carried out with the NucleoSpin RNA virus kit (Macherey-Nagel, 740956) as described in the manufacturers’ instructions. In order to analyze the presence of FL vRNA and DI vRNA (truncated form), a segment-specific RT-PCR was performed (see 4.4.1). Real-time RT-qPCR was applied for absolute quantification of Seg 5 vRNA from the progeny virions (see 4.4.2).

#### 4.4.1 Segment-specific RT-PCR

A recently described method was utilized for segment-specific RT-PCR (17, 30). In brief, isolated vRNA was reverse transcribed into cDNA using universal primers that bind at the conserved terminal regions of all eight IAV genome segments (Supplementary Table 1). Subsequently, segment-specific primers were used for amplification of the respective genome segment sequence by PCR (Supplementary Table 1). Finally, PCR products were analyzed by agarose gel electrophoresis.

#### 4.4.2 Real-time RT-qPCR

A recently reported method for the specific detection and quantification of influenza viral RNA segments using real-time RT-qPCR was employed (17, 57, 58). Briefly, RNA reference standards were *in vitro* synthesized for absolute quantification (primers required for generation are listed in Supplementary Table 4). Isolated vRNA of the samples was used for reverse transcription, along with a dilution series of the reference standards (primers listed in Supplementary Table 2), followed by real-time qPCR (primer sequence in Supplementary Table 3). Calculation for absolute quantification of vRNA of Seg 5 was conducted as previously described (17, 58).

#### 4.4.3 NGS and data processing

Sample preparation, NGS library preparation and sequencing analysis of deletion junctions was performed according to a recently published study (32).

### 4.5 Analysis of deletion junctions

Deletion junctions refer to the DI vRNAs in the viral population, while deletion junction sites refer to the start and end position of the breaking points in the viral genome. Deletion junctions that did not accumulate to levels above 14 NGS reads in at least one sampling time point were removed from the data set for higher accuracy (32). Furthermore, defective vRNAs that showed more than 85 % of the length of FL vRNA were excluded from analysis in this work (except for Supplementary Fig. 3). DI vRNA 3’ and 5’ length indicated the number of retained nucleotides prior and after the deletion junction at the respective vRNA end. Of note is that DI vRNA sequence was reported in negative-sense and 3’ to 5’ orientation. The calculation of the DI vRNA length comprised the following sequence lengths: Seg 1 (2341 nucleotides (nt)), Seg 2 (2341 nt), Seg 3 (2233 nt), Seg 4 (1775 nt), Seg 5 (1565 nt), Seg 6 (1413 nt), Seg 7 (1027 nt), Seg 8 (890 nt). Number of nucleotides of the incorporation signal (NCR) and the bundling signal (terminal ends of coding region), which together form the packaging signal, were taken from a recent review (59).

### 4.6 Generation of purely clonal DIPs containing a deletion in Seg 1

To generate purely clonal Seg 1 derived DIPs (top gain, loss, gain (*de novo*)) in the absence of STV, we used a previously established plasmid-based reverse-genetics system (41). More specifically, to complement the missing PB2 to allow DIP production without STV, we used a co-culture of HEK-293T-PB2 cells and MDCK-PB2(adh) cells for reconstitution.

#### 4.6.1 Generation of plasmids

Plasmids harboring specific deletions were generated as described previously (41). In brief, pHW191 encoding the PR8-derived PB2 gene (60) was used as a template for PCR amplification (Phusion Hot Start II DNA polymerase, Thermo Fisher, #F549L). Here, the desired 5’-fragment (containing overhangs complementary to the 3’-fragment) of a specified deletion junction (Supplementary Table 5), using a 5’-specific forward and reverse primers set (Supplementary Table 6) was used. Similarly, a set of 3’-specific primers were used to amplify the desired 3’-fragment (containing overhangs complementary to the 5’-fragment) of a specified deletion junction from the pHW191 template DNA. Next, the 5’-fragments hybridized with the overlapping 3’-fragments, resulting in PCR products with the individual deletion junctions (splice-overlapped products) after subsequent amplification cycles at an annealing temperature of 62°C. Lastly, the internally spliced PB2 sequence was inserted in pHW2000-GGAarI using golden-gate cloning (61, 62). All plasmids were sequenced to confirm the generated deletion junctions.

#### 4.6.2 Rescue and production of DIPs

For rescue of purely clonal DIPs containing a deletion in Seg 1 (41), we co-transfected a co-culture of adherent HEK-293T-PB2 cells (0.2 × 10^6^ cells/well) and MDCK-PB2(adh) cells (0.2 × 10^6^ cells/well) with corresponding plasmids harboring a deletion in the PB2 sequence (50 ng) and 1 μg of each pHW192-pHW198 plasmid (encoding for the remaining gene segments of PR8 IAV) using the calcium phosphate method in a 6-well format. DIP-containing supernatants were harvested at 4, 6, 8, 10, and 12 days post transfection and stored at -80°C for further use. Larger stocks (seed viruses) of purely clonal Seg 1 DIPs were generated in MDCK-PB2(sus) cells in shake flasks.

The production of Seg 1 DIPs in MDCK-PB2(sus) cells was conducted according to a recently published paper (25). In brief, MDCK-PB2(sus) cells, cultivated in shake flasks were centrifuged (300×g, 5 min, room temperature) and used to inoculate a new shake flask at 2.0 × 10^6^ cells/mL with fresh media and trypsin (final activity of 20 U/mL). Subsequently, cells were infected at an MODIP of E-2. Cell-free supernatants (3000×g, 4°C, 10 min) were stored at -80°C for further analysis.

### 4.7 Interference assay

To measure the efficacy of DIPs to suppress STV replication, we performed an *in vitro* co-infection assay in MDCK(adh) cells following a previously published description (26). To summarize, MDCK(adh) cells, cultivated in 6-well plates, were washed twice with phosphate-buffered saline (PBS). Next, cells were either infected with STV only (MOI 0.01, based on the TCID_50_ titer) or co-infected with STV and 125 µL of the produced DIP material (diluted for normalization, Supplementary Table 7). Wells were filled up to 250 µL with infection medium (GMEM, 1% peptone, 5 U/mL trypsin) and incubation was conducted for 1 h at 37°C and 5% CO_2_. Subsequently, the inoculum was aspirated, the cells washed with PBS and 2 mL of infection medium was added. Cells were incubated for 24 h at 37°C and 5% CO_2_. The supernatant was harvested and stored at -80°C until further analysis by plaque assay and HA assay.

### 4.8 Data availability

The reference sequence of the PR8 genome used for alignment can be found under the following NCBI accession numbers: PB2 = AF389115.1, PB1 = AF389116.1, PA = AF389117.1, HA = AF389118.1, NP = AF389119.1, NA = AF389120.1, M = AF389121.1, NS = AF389122.1. The complete NGS dataset is available under the BioProject accession number PRJNA743179.

## 6 Author Contributions

Conceptualization, L.P., D.R., T.D., Y.G., C.B.B., S.Y.K. and U.R.; Formal analysis, L.P., D.R., F.G.A and S.Y.K.; Funding acquisition, C.B.B. and U.R; Investigation, L.P., D.R., T.D. and F.G.A.; Project administration, L.P. and S.Y.K.; Supervision, Y.G., C.B.B., S.Y.K. and U.R.; Visualization, L.P., D.R. and S.Y.K.; Writing – original draft, L.P., T.D. and S.Y.K.; Writing – review & editing, L.P., D.R., T.D., F.G.A., Y.G., C.B.B., S.Y.K. and U.R..

## 7 Acknowledgments

Special thanks to Claudia Best and Nancy Wynserski for their outstanding technical support. We thank Thomas Bissinger for providing the seed virus and Marc Hein for valuable discussions. The authors appreciate the sequencing work of Alvaro Hernandez, Christy Wright and the staff at the Roy. J. Carver Biotechnology Center, University of Illinois. We would like to thank St. Jude children research hospital, Memphis, United States for providing pHW2000 plasmids. Furthermore, a special thank is directed to Prof. Stefan Pöhlmann, Prerna Arora, and Najat Bdeir of the German Primate Center, Göttingen, Germany for providing plasmids and cell lines for rescue of DIP candidates and helpful discussions. We thank Shanghai BioEngine Sci-Tech and Prof. Tan from the East China University of Science and Technology for supplying the Xeno™ medium. The work was supported by the Defense Advanced Research Projects Agency (https://www.darpa.mil/program/intercept), INTERCEPT program under Cooperative Agreement W911NF-17-2-0012 and DARPA-16-35-INTERCEPT-FP-018.

