## Supplementary Material for "Semi-continuous propagation of influenza A virus and its defective interfering particles: analyzing the dynamic competition to select candidates for antiviral therapy"

### 1 Supplementary Material

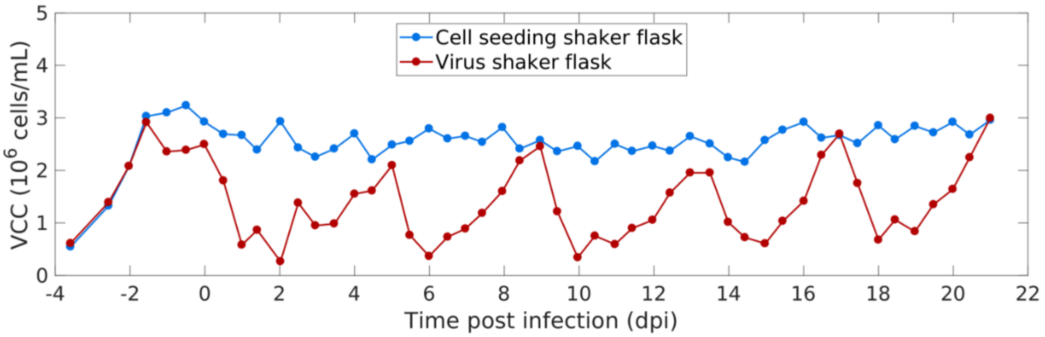


**Supplementary Figure 1: Viable cell concentration during semi-continuous production of IAV.** MDCK(sus) cells were grown in the cell seeding shake (CSS) flask and the virus shake (VS) flask. After an initial batch and semi-continuous cultivation phase with uncoupled CSS and VS, cells in the VS were infected with the seed virus at an MOI of 0.1. The semi-continuous production mode was initiated 0.5 dpi, where cells were transferred from the CSS into the VS at regular time intervals (12 h), while fresh medium was added and virus harvest was taken for analytics. Illustration includes results of one experiment.

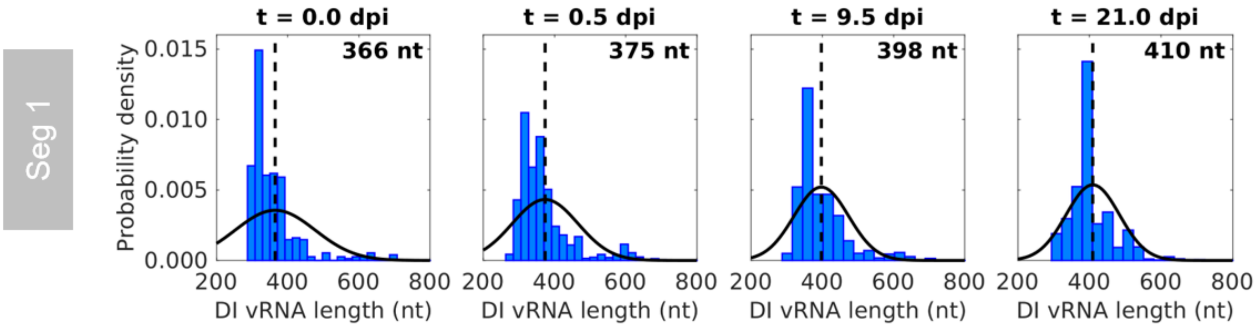


**Supplementary Figure 2: Normal distributions and means of DI vRNA length.** Deletion junctions were identified by Illumina-based NGS and subsequently analyzed via the ViReMa algorithm (32). Means of the DI vRNA length based on normal distribution functions fitted (indicated by a dashed black line). Corresponding lengths are indicated, Seg 1 is shown as an example. Representative time points are illustrated. Illustration includes results of one experiment.

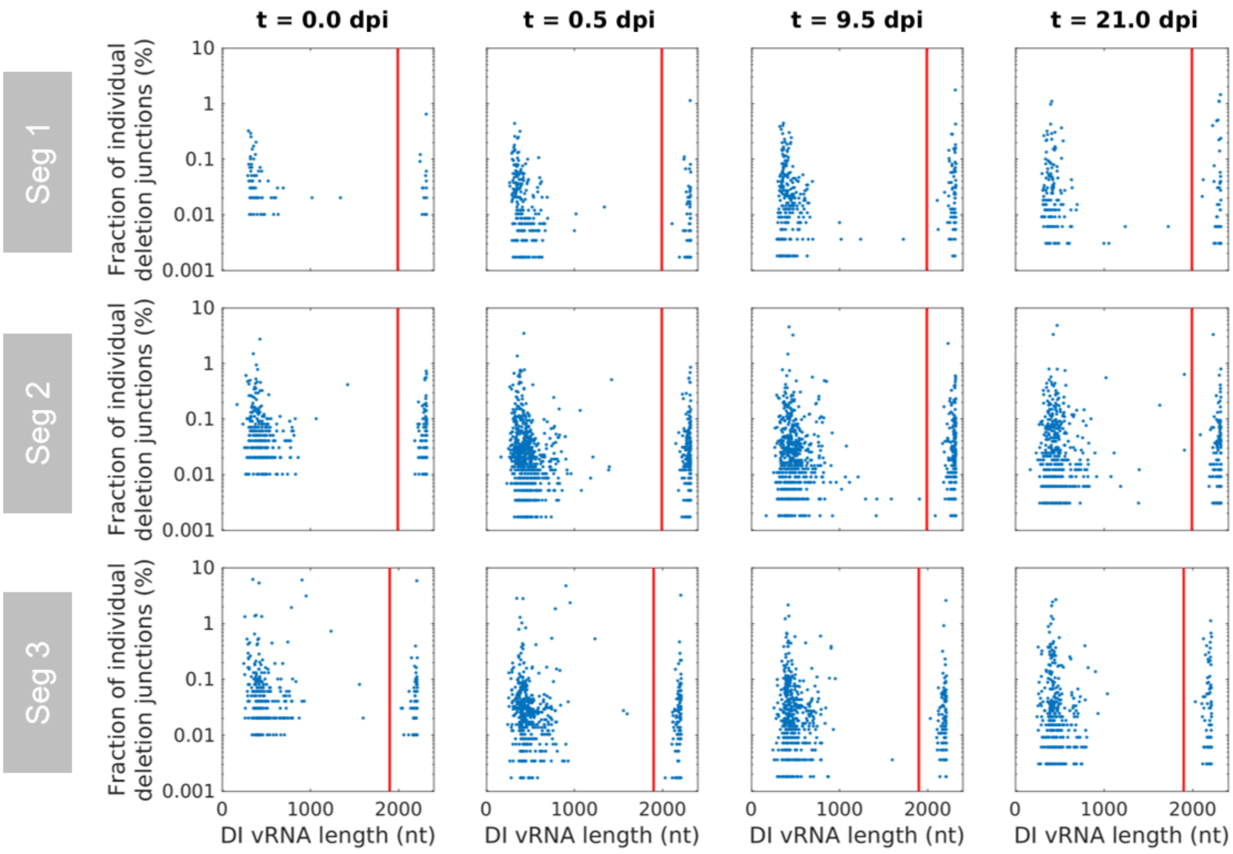


**Supplementary Figure 3: DI vRNAs up to a maximum length of 2400 nt.** Deletion junctions were identified by Illumina-based NGS and subsequently analyzed via the ViReMa algorithm (32). Fractions of individual deletion junctions were calculated based on the ratio of the number of NGS reads of one individual deletion junction to the number of NGS reads of all deletion junctions located on all eight segments. Very long DI vRNAs (indicating very short deletions) detected by NGS and excluded for analysis. Vertical red line indicates the cut-off used for excluding these vRNAs (85% of FL length). Representative time points are illustrated. Illustration includes results of one experiment.

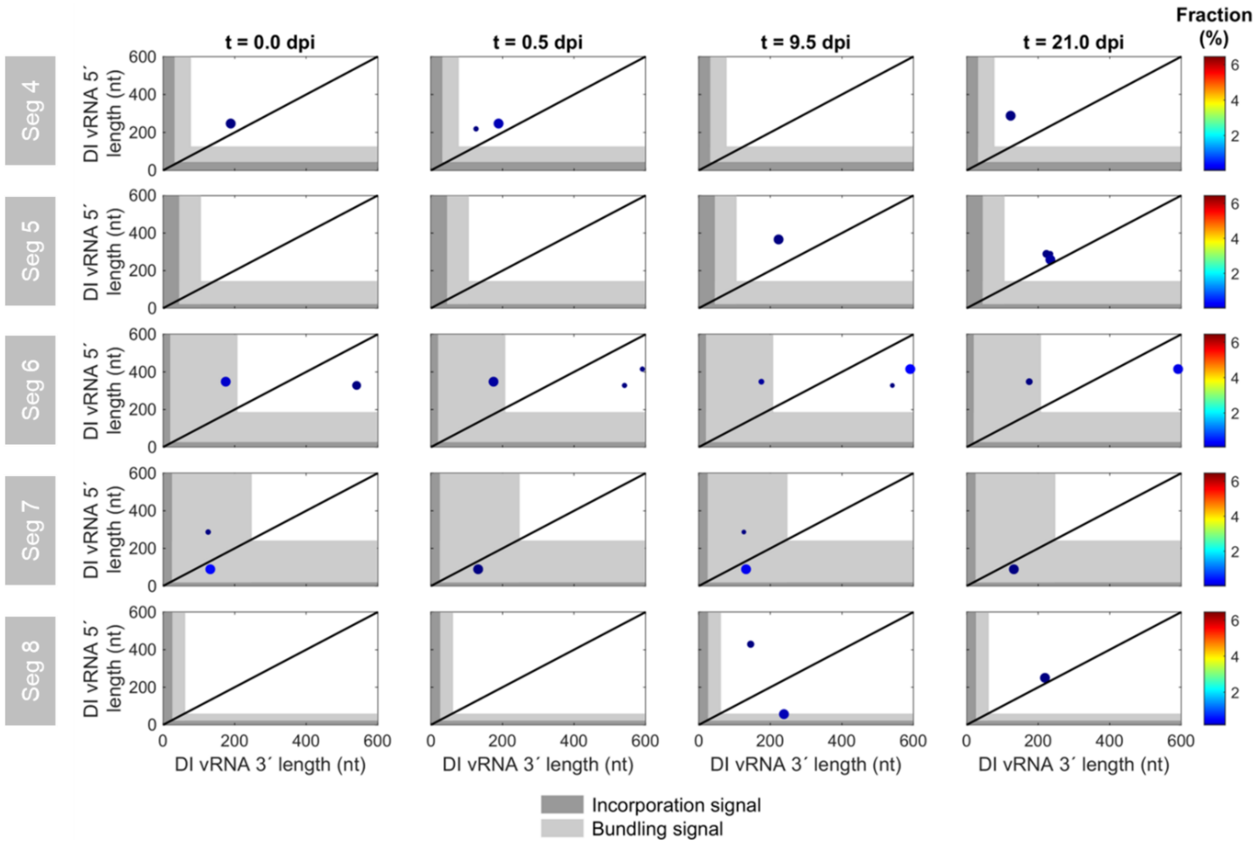


**Supplementary Figure 4: Position of DI vRNA breaking points on non-polymerase-encoding segments during semi-continuous propagation of IAV.** Deletion junctions were identified by Illumina-based NGS and subsequently analyzed via the ViReMa algorithm (32). DI vRNA 3´ and 5' length indicate the number of retained nucleotides prior and after the deletion junction at the respective vRNA ends. The packaging signal is indicated as grey area, which is divided into the incorporation signal (dark grey area) and bundling signal (light grey area). Representative time points are shown. The color code from red to blue shown on the right denotes the fraction of individual deletion junction, which was calculated based on the ratio of the number of NGS reads of one individual deletion junction to the number of NGS reads of all deletion junctions located on all eight segments.For graphs where no breaking points are shown, no DI vRNAs were detected at the selected time points. The diagonal black line indicates an equal DI vRNA 3´ and 5´length. Illustration includes results of one experiment.

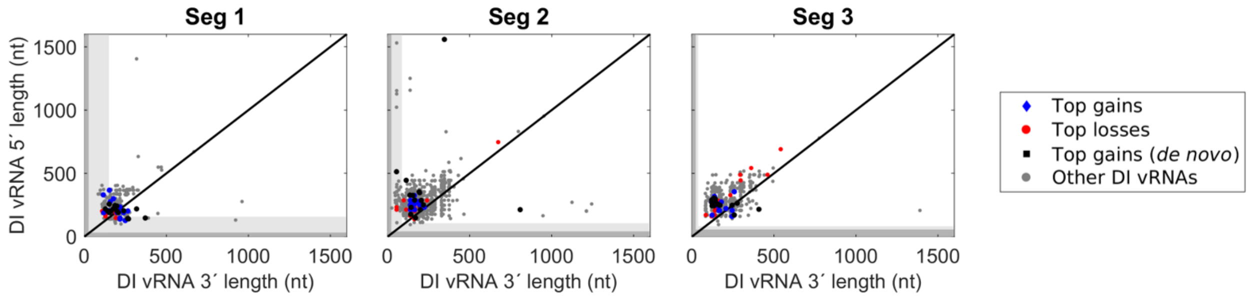


**Supplementary Figure 5: Deletion junctions with the highest increase or decrease in the fraction during semi-continuous propagation of IAV.** Deletion junctions were identified by Illumina-based NGS and subsequently analyzed via the ViReMa algorithm (32). DI vRNA 3´ and 5’ length indicate the number of retained nucleotides prior and subsequent to the deletion junction at the respective vRNA ends. The packaging signal is indicated as grey area, which is divided into the incorporation signal (dark grey area) and bundling signal (light grey area). Breaking points of the top 15 gains up to a DI vRNA length of 1600 nt. On Seg 2, two top 15 gains (*de novo*) candidates were found comprising a long DI vRNA length.

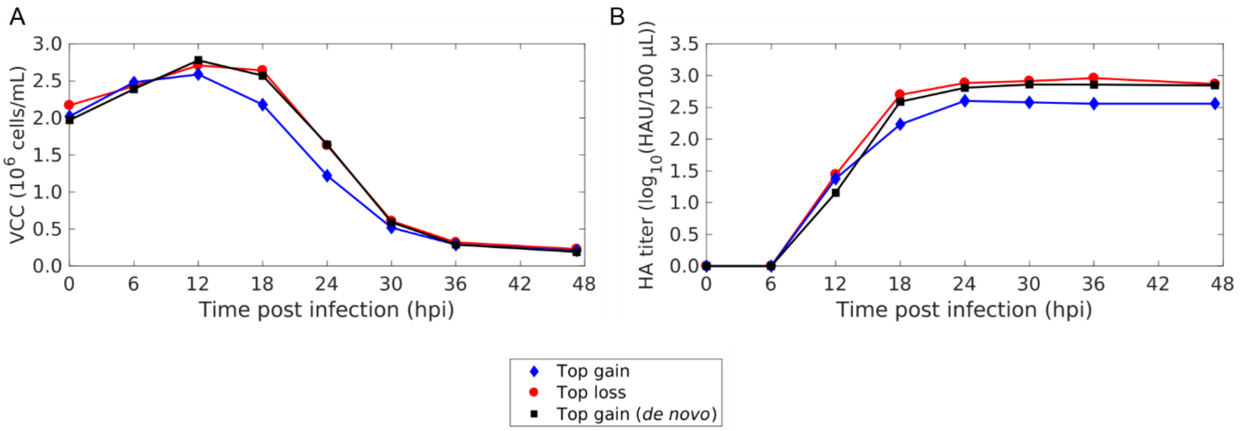


**Supplementary Figure 6: Batch production of Seg 1 DIPs derived from top gain, loss, and gain (*de novo*) DI vRNAs.** MDCK-PB2(sus) cells were seeded in shake flask at $2.0 \times 10^6$ cells/mL after a full medium exchange. Infection with DIPs was carried out at MODIP E-2. (A) VCC. (B) HA titer. Illustration includes results of one experiment.

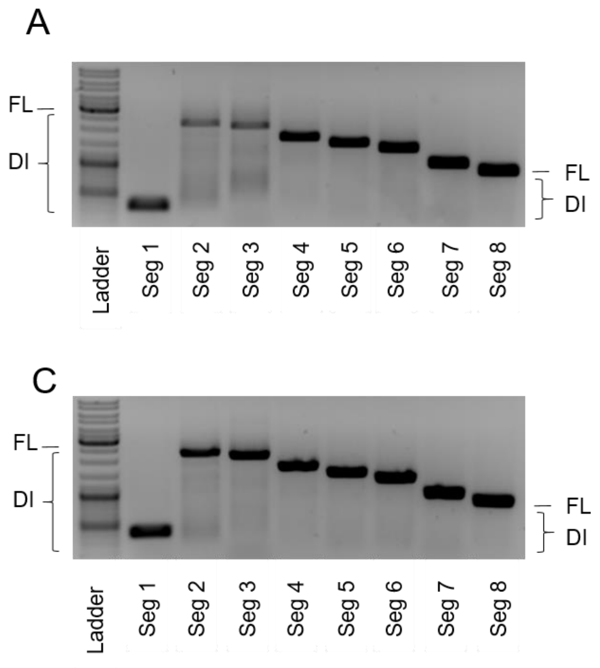


**Supplementary Figure 7: Integrity of Seg 1 DIPs of the batch production.** Results of the segment-specific RT-PCR for Seg 1 to 8 followed by agarose gel electrophoresis are shown. (A) Top gain. (B) Top loss. (C) Top gain (*de novo).* Corresponding lengths are in agreement with the expected size (Supplementary Table 5). Only weak, if any, additional bands occurred. Signals corresponding to FL and DI vRNAs are indicated. Upper, middle and lower thick bands of the DNA ladder indicate 3000, 1000 and 500 bp, respectively.

**Supplementary Table 1: Primers used for segment-specific RT-PCR.**

| **Reaction** | **Target** | **Primer name** | **Sequence (5´→3´)** |
|---|---|---|---|
| **RT** | All segments | Uni 12 | AGCAAAAGCAGG |
| **PCR** | Segment 1 | S1 Uni for | AGCGAAAGCAGGTCAATTAT |
| | | S1 Uni rev | AGTAGAAACAAGGTCGTTTTTAAAC |
| | Segment 2 | S2 Uni for | AGCGAAAGCAGGCAAACCAT |
| | | S2 Uni rev | AGTAGGAACAAGGCATTTTTTCATG |
| | Segment 3 | S3 Uni for | AGCGAAAGCAGGTACTGATCC |
| | | S3 Uni rev | AGTAGAAACAAGGTACTTTTTTGG |
| | Segment 4 | S4 Uni for | AGCAAAAGCAGGGGAA |
| | | S4 Uni rev | AGTAGAAACAAGGGTGTTTT |
| | Segment 5 | S5 Uni for | AGCAAAAGCAGGGTAGATAATC |
| | | S5 Uni rev | AGTAGAAACAAGGGTATTTTTC |
| | Segment 6 | S6 Uni for | AGCGAAAGCAGGGGTTTAAAATG |
| | | S6 Uni rev | AGTAGAAACAAGGAGTTTTTTGAAC |
| | Segment 7 | S7 Uni for | AGCGAAAGCAGGTAGATATTG |
| | | S7 Uni rev | AGTAGAAACAAGGTAGTTTTTTAC |
| | Segment 8 | S8 Uni for | AGAAAAAGCAGGGTGACAAA |
| | | S8 Uni rev | AGTAGAAACAAGGGTGTTTT |

**Supplementary Table 2: Primers used for RT.**

| Target | Primer name | Sequence (5´→3´) |
|---|---|---|
| Seg 5 | S5 tagRT for | ATTTAGGTGACACTATAGAAGCGAGTGATTATGAGGGACGGTTGAT |

**Supplementary Table 3: Primers used for real-time qPCR.**

| Target | Primer name | Sequence (5´→3´) |
|---|---|---|
| Introduced tag sequence | vRNA tagRealtime for | ATTTAGGTGACACTATAGAAGCG |
| Seg 5 | Seg 5 Realtime rev | CGCACTGGGATGTTCTTC |

**Supplementary Table 4: Primers used for reference standard generation.**

| Target | Primer name | Sequence (5´→3´) |
|---|---|---|
| Seg 5 | S5 uni for | AGCAAAAGCAGGGTAGATAATC |
|  | S5 uni T7 rev | TAATACGACTCACTATAGGGAGTAGAAACAAGGGTATTTTTC |

**Supplementary Table 5: Generated Seg 1 candidate DIPs, and the respective deletion junction positions in the 5´-3´ cDNA sequence.**

| DIP | 5’ fragment size | 3’fragment size | DIP fragment size |
|---|---|---|---|
| Loss | 129 bp | 166 bp | 295 bp |
| Gain | 217 bp | 138 bp | 355 bp |
| Gain (*De novo*) | 269 bp | 140 bp | 409 bp |

**Supplementary Table 6: Splice overlap extension PCR primers.**

| Target | Primer name | Sequence (5´→3´) | Annealing temperature for overlap PCR |
|---|---|---|---|
| Gain 5' fragment | For | CGGTCACCTGCCAGTGGGAGCGAAAGCA | 66°C |
| | Rev | ACGTCTCCTTGCCCAATTATCCTCTTGTCTGCTGTA | |
| Gain 3' fragment | For | TACAGCAGACAAGAGGATAATTGGGCAAGGAGACGT | |
| | Rev | CCCACCTGCGCGCTATTAGTAGAAACAAGGTCGTTTTTAAACTATT | |
| Loss 5' fragment | For | CCCACCTGCCAGTGGGAGCGAAAGCAG | 67°C |
| | Rev | GCCTTCTCTCCTTTCGCGTACTTCTTGATTATGGCCA | |
| Loss 3' fragment | For | TGGCCATAATCAAGAAGTACGCGAAAGGAGAGAAGGC | |
| | Rev | CCCACCTGCGCGCTATTAGTAGAAACAAGGTCGTTTTTAAACTA | |
| Gain (*de novo*) 5' fragment | For | CAGTCACCTGCCGATGGGAGCGAAAGCAGGT | 67°C |
| | Rev | CCACGTCTCCTTGCCCAATTATTTTACTCCATAAAGTTTGTCCTTGC | |
| Gain (*de novo*) 3' fragment | For | GCAAGGACAAACTTTATGGAGTAAAATAATTGGGCAAGGAGACGTGG | |
| | Rev | CCCACCTGCTTTTTATTAGTAGAAACAAGGTCGTTTTTAAACTATTC | |

**Supplementary Table 7: DIPs and STVs used for interference assay.** Production titers are indicated. DIP input was normalized through dilution based on the concentration of DIPs derived from HA titer. DI244 was chosen as a control.

| **Sample** | **HA titer** log10(HAU/100µL) | **Concentration of DIPs** DIPs/mL | **Dilution factor** | **Concentration of STVs** STVs/mL |
|---|---|---|---|---|
| **Gain** | 2.23 | $3.43 \times 10^9$ | 1.00 | - |
| **Loss** | 2.70 | $9.97 \times 10^9$ | 2.91 | - |
| **Gain *(de novo)*** | 2.59 | $7.74 \times 10^9$ | 2.26 | - |
| **DI244** | 2.58 | $7.56 \times 10^9$ | 2.20 | - |
| **STV** | 2.72 | - | - | $1.05 \times 10^{10}$ |
